# Lysosomal abundance in young and aged mouse hearts assessed by In Vivo Imaging Systems (IVIS) Lysotracker imaging and autophagy-related gene expression

**DOI:** 10.64898/2026.02.16.706145

**Authors:** Jawaher Albulushi, Hannah Coghlan, Mohesh Moothanchery, Aiswarya Dev, Emily Akerman, Jasmine Heenan, Nordine Helassa, Oluwatobi Adegbite, Parveen Sharma, Fenil Patel, Libby Harrison, Mahon L Maguire, Gary R Mirams, Pawel Sweitach, Harish Poptani, Rebecca AB Burton

## Abstract

Lysosomal function is essential for cardiac proteostasis and cellular health, yet its regulation during ageing remains poorly defined. We hypothesised that ageing alters both the abundance of acidic organelles and the machinery supporting their acidification. Using fluorescence-based In Vivo Imaging Systems (IVIS) with Lysotracker™ Red in young (2–4 months) and aged (18 months) mouse hearts, we quantified whole-heart acidic-vesicle signals and assessed expression of lysosomal and autophagy-related genes (*Lamp2, Atp6v1a, Sqstm1, Cd63, Atg12, Nfe2l2, M6pr*) by RT-qPCR. Whole-heart labelled Lysotracker fluorescence did not differ significantly between age groups, indicating preservation of the total acidic-vesicle pool. No changes in *Atp6v1a* and *Lamp2* expression suggest acidification capacity and structural stability are maintained, whereas the minor, upregulation of *Sqstm1* might indicate increased autophagic demand and altered vesicle trafficking, which warrants further investigation. No statistical significant changes in *M6pr, Atg12, or Nfe2l2* were detected, suggesting transcriptional stability in enzyme trafficking, core autophagy, and oxidative stress pathways. Regionally, atria showed higher Lysotracker signal than ventricles, consistent with known enrichment of acidic vesicular stores in atrial physiology. These findings highlight the utility of IVIS imaging of Lysotracker-labelled hearts, providing rapid whole-organ assessment of acidic vesicle distribution, albeit with limited depth resolution. Complementary techniques such as RT-qPCR analysis is essential to interpret IVIS findings, enabling insight into underlying molecular changes in lysosomal and autophagy pathways during cardiac ageing.

## Introduction

Ageing is an inevitable biological process and the most significant non-modifiable risk factor for cardiovascular disease (CVD)(*1*). As the population ages, the burden of CVD is projected to rise markedly. Between 2025 and 2050, cardiovascular deaths are expected to increase from 20.5 million to 35.6 million, a 73.4% rise, primarily due to demographic ageing (*2*). Although age-standardized mortality rates are anticipated to decline by approximately 30.5% due to improvements in diagnosis and treatment, the absolute number of cases and deaths will continue to grow, underscoring the need for targeted healthcare strategies in ageing populations (*2*).

In humans, the ageing heart undergoes both structural and functional changes, including thickening of the left ventricle (LV), diastolic dysfunction, fibrosis, and reduced cardiac output under stress (*1, 3*). Even without comorbidities such as hypertension or diabetes, these intrinsic alterations elevate mortality risk by diminishing cardiac efficiency and adaptability (*3*). At the cellular level, ageing is characterized by elevated oxidative stress, mitochondrial dysfunction, reduced metabolic flexibility, and increased cardiomyocyte senescence (*1, 4*). These changes compromise energy homeostasis and calcium (Ca^2+^) handling, contributing to progressive cardiac impairment. Increasing attention has focused on the failure of cellular quality control systems, particularly autophagy and lysosomal degradation, which are essential for clearing damaged proteins and organelles (*4, 5*).

Historically regarded as passive sites of cellular waste disposal, lysosomes were first identified in 1955 by Christian de Duve through subcellular fractionation (*6*), a discovery that laid the foundation for modern cell biology (*7*). They are now recognized as dynamic organelles with critical roles in metabolic regulation, intracellular signalling, and inter-organelle communication (*8*). This evolving understanding has positioned lysosomes as central players in the pathophysiology of ageing and chronic diseases (*8*).

Within cardiomyocytes, lysosomes have emerged as key regulators of Ca^2+^ homeostasis, energy metabolism, and cellular stress responses (*9, 10*). A growing body of research has identified nicotinic acid adenine dinucleotide phosphate (NAADP) as a key signalling molecule that specifically targets lysosomes (*11*). NAADP activates two-pore channels (TPCs) on the lysosomal membrane, triggering the release of lysosomal Ca^2+^ (*12*). TPCs act as NAADP receptors, releasing Ca^2+^ from acidic organelles, which can then initiate additional Ca^2+^ signalling through the sarcoplasmic or endoplasmic reticulum (*12*). Notably, this pathway operates independently of the traditional Ca^2+^ influx through L-type calcium channels (*9*).

Further studies show that lysosomal Ca^2+^ stores are essential for modulating how heart cells respond to stress signals like β-adrenergic stimulation (*9*). Among the TPC family, TPC2 appears to be the dominant channel involved in generating these NAADP-triggered Ca^2+^ signals (*9*). The use of pharmacological agents such as bafilomycin A1 and glycyl-L-phenylalanine 2-naphthylamide (GPN) has provided functional evidence that NAADP-evoked Ca^2+^ signals originate specifically from lysosomes (*13*).

Additionally, during ischemia–reperfusion injury (a type of damage that occurs when blood returns to the heart after a period of oxygen deprivation), NAADP-mediated signalling triggers Ca^2+^ release from lysosomes. This lysosomal Ca^2+^ release promotes pathological Ca^2+^ oscillations that drive mitochondrial Ca^2+^ overload and ultimately cardiomyocyte death (*14*). Pharmacological inhibition of NAADP signalling at the onset of reperfusion significantly reduced infarct size and improved cardiac outcomes in experimental models, implicating lysosomal Ca^2+^ release as an early initiator of reperfusion-induced injury (*14*).

Beyond their role in Ca^2+^ signalling, lysosomes are essential regulators of cellular energy balance and protein quality control. They closely interact with mitochondria and influence key pathways such as autophagy and mTOR signalling, both of which are frequently disrupted during cardiac ageing and disease (*1, 15*). Impaired lysosomal acidification or blocked autophagic flux can trigger pathological conditions, including myocardial infarction and cardiac hypertrophy (*1, 15*). In the ageing myocardium, particularly in long-lived, non-dividing cardiomyocytes, lysosomal degradative capacity declines, leading to the buildup of oxidized proteins, dysfunctional organelles, and lipofuscin. This overload compromises mitochondrial recycling and reduces the heart’s contractile function and stress resilience over time (*4, 5, 15*).

Atrial fibrillation (AF) is one of the most common arrhythmias, and its prevalence increases markedly with ageing. Endolysosomal dysregulation as well as lysosomal special reorganisation of lysosomes (*10, 16*) has been observed in experimental models of AF and patients with chronic AF, suggesting a potential mechanistic link between ageing-related cellular changes and arrhythmogenesis (*17*). This suggests that lysosomes and related acidic compartments may play a role in the structural and metabolic remodelling seen in AF (*17*). These findings support the idea that lysosomes serve as both sensors and effectors of cardiac stress, that might drive disease progression (*8*).

Recent research has substantially advanced our understanding of lysosomal dysfunction in acute cardiac conditions (*18*), such as myocardial ischemia-reperfusion injury (MI/RI), diabetic cardiomyopathy (DCM), and lysosomal storage disorders (LSDs). However, opportunities remain to deepen our knowledge of lysosomal involvement in ageing-associated chronic cardiac remodelling. While most studies have focused on the acute stress response, emerging evidence highlights the importance of understanding lysosomal regulation in the ageing heart and its contribution to progressive myocardial decline (*19-21*). Promising strategies such as the transcription factor EB (TFEB) activation (a master regulator of lysosomal biogenesis and autophagy), autophagy modulation, and gene therapy have shown considerable potential for restoring lysosomal function (*20, 22, 23*). Expanding these approaches to age-related cardiomyopathies represents an exciting avenue for future research, with the possibility of developing targeted, lysosome-based therapeutic interventions.

Among emerging therapeutic strategies, targeting lysosomal Ca^2+^ signalling has gained particular attention. Modulation of TPC2 channels has demonstrated anti-arrhythmic effects in models of catecholaminergic stress by suppressing aberrant Ca^2+^ release events (*9*). While these findings highlight the potential of lysosome-based interventions for cardiac electrophysiology, their physiological relevance during age-related structural and metabolic remodelling remains to be fully understood. In parallel, combinatorial approaches integrating TFEB activators, autophagy modulators, and gene therapies are showing increasing promise; however, their application to the ageing myocardium remains in the early experimental stages (*23*).

Imaging technologies have advanced lysosomal research, with tools like fluorescence microscopy and *in vivo* imaging systems (like IVIS) offering high-resolution, real-time insights into lysosomal dynamics (*24-26*). However, optical imaging remains limited by shallow tissue penetration and signal scattering in deep cardiac tissues (*25*). Non-invasive modalities such as magnetic resonance imaging (MRI), positron emission tomography (PET), and computed tomography (CT) provide whole-organ views, MRI excels in soft-tissue contrast, PET in metabolic sensitivity, and CT in anatomical resolution, including microscopic detail in some settings (*24, 25*). Yet, these medical imaging technologies lack the molecular specificity to directly assess lysosomal compartments. Techniques like chemical exchange saturation transfer (CEST) MRI show promise for tracking lysosomal pH but are limited by motion artifacts and the absence of targeted contrast agents (*26*). These gaps highlight the need for integrative, multimodal strategies to achieve selective, *in vivo* assessment of lysosomal function in the ageing heart.

In this study, we hypothesised that ageing would alter the abundance of acidic organelles and their acidification machinery in cardiac tissue. To test this, we used IVIS fluorescence imaging of Lysotracker to quantify whole-heart Lysotracker uptake as a proxy for the pool of acidic vesicles (*27*), and real-time polymerase chain reaction (RT-qPCR) to assess key lysosomal genes, including *Lamp2* (lysosomal structure), *Cd63* (late endosome/multivesicular body (MVB) marker), *M6pr* (endosomal trafficking), *Atg1*2 (autophagy initiation), *Sqstm1* (autophagy adaptor), *Nfe2l2* (oxidative stress regulator), and *Atp6v1a* (a v-ATPase subunit essential for lysosomal acidification) (*28*)).

## Results

To assess whether lysosomal abundance changes with age in the heart, we compared Lysotracker fluorescence in *ex vivo* hearts from young and old mice (3 animal batches, n = 7 per age group) using IVIS imaging. Quantitative analysis of total cardiac fluorescence revealed no significant difference between age groups (*p* > 0.05). Given the surface-weighted nature of optical imaging, these measurements could primarily reflect lysosomal signal from superficial myocardial layers and should be interpreted as semi-quantitative. Thus, while our data suggest comparable epicardial lysosomal activity between age groups, we cannot exclude age-related changes in deeper myocardial regions that may be underestimated due to depth-dependent attenuation. Additionally, age-associated alterations in myocardial structure or optical properties could influence fluorescence detection independently of lysosomal biology. Together, these findings indicate that global myocardial Lysotracker signal appears preserved with aging under our imaging conditions, though complementary *ex vivo* or histological analyses will be required to fully assess potential transmural or cell-type–specific differences. To complement these findings, we measured the expression of the lysosomal membrane protein gene *Lamp2* by RT-qPCR in whole-heart RNA extracts. Consistent with the imaging results, *Lamp2* transcript levels did not differ significantly between young and aged hearts. Together, these data suggest that gross lysosomal abundance is maintained in the aging heart, although subtle or region-specific alterations may not be captured by surface-weighted imaging or bulk transcript analysis.

### IVIS Imaging

Animals were purchased in three batches (see Methods). Visual inspection of the raw IVIS data (Figure 1) showed that atrial signals were consistently higher than ventricular signals in both age groups. In batch 2, old mice appeared to have lower signals than young mice. Overall signal intensity also varied between batches.

**Figure 1.**
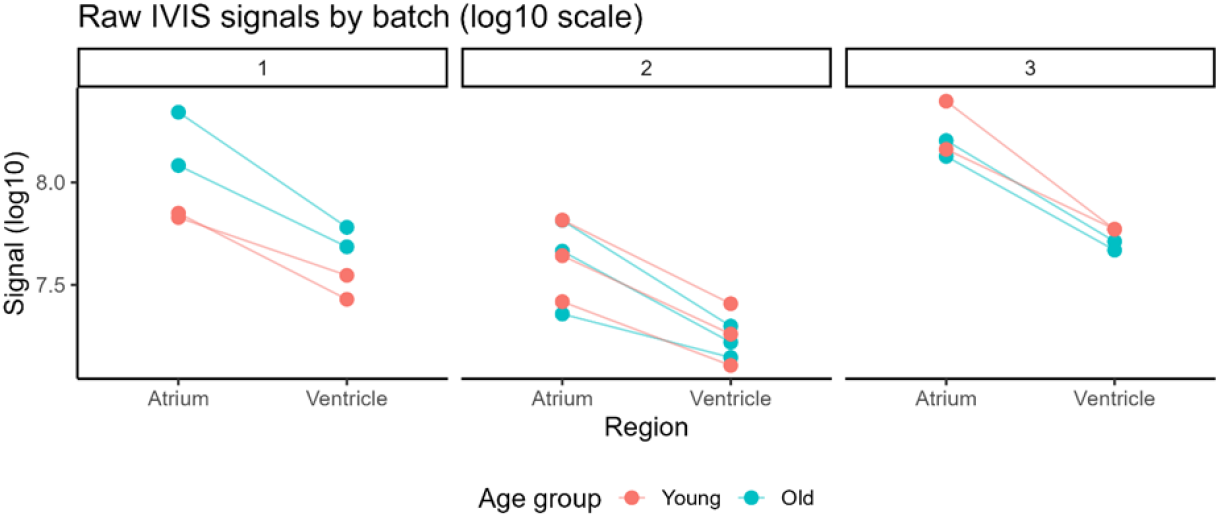
Raw Lysotracker fluorescence IVIS signals by batch (log10 scale). Raw IVIS signals (log10 scale) for atrium and ventricle in young and old mice, shown separately for each imaging batch. Lines connect atrium and ventricle values from the same mouse. In some batches, old mice appeared to have slightly higher or lower signals than young mice, but these differences were inconsistent in direction across batches. Old animals n=7, young animals n=7.

Given these patterns, and the paired nature of the data (atrium and ventricle from the same mouse), a linear mixed-effects model was used (see Supplementary Table S1 for full estimated marginal means (EMMs), Supplementary Figure S1 for diagnostic plots (i.e. model-checking plots assessing assumptions of the linear mixed-effects model), and Supplementary Table S2 for the full ANOVA results), with Age group, Region, and Batch as fixed effects, and Mouse ID as a random effect. The analysis confirmed a significant effect of Region (F(1,13) = 207.31, *p* < 0.0001) and Batch (F(2,10) = 17.30, *p* = 0.0006), but no significant effect of Age group (F(1,10) = 0.38, *p* = 0.553). Batch-adjusted means (Figure 2) indicated that, on the log10 scale, atrial signals averaged between 7.93–7.98 for young and old mice, whereas ventricular signals averaged between 7.51–7.56. Back-transformed geometric means correspond to ∼8.45–9.48 × 10^7^ photons/sec/cm^2^/sr for atria and ∼3.21–3.60 × 10^7^ for ventricles, with overlapping 95 % confidence intervals between age groups.

**Figure 2.**
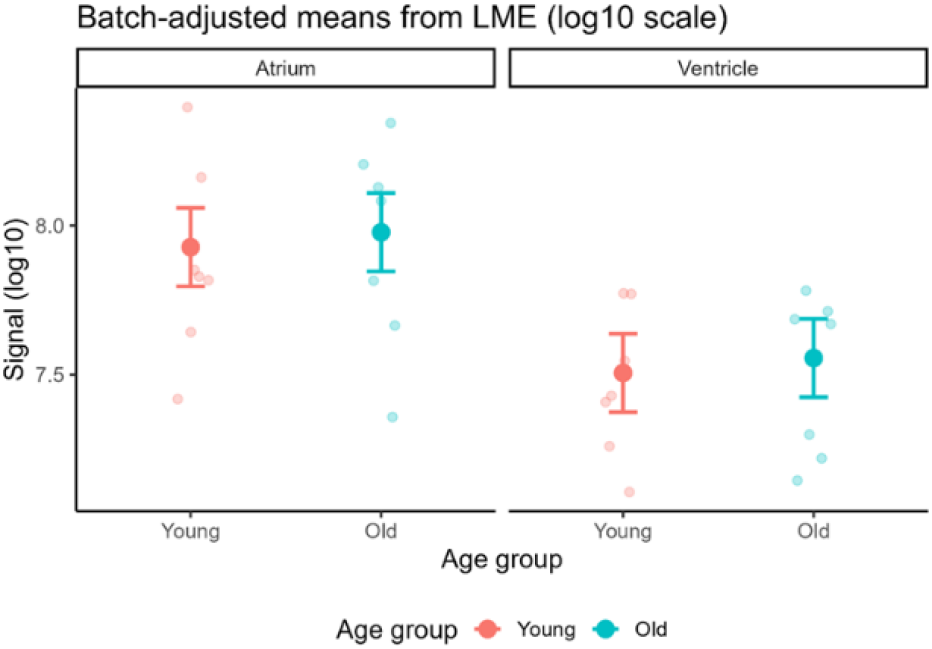
Batch-adjusted means (log10 scale) from the linear mixed-effects model for atrium and ventricle in young and old mice. Points represent estimated marginal means (±95 % CI) after adjusting for batch effects; faint points show individual mouse values. Batch-adjusted means indicate higher atrial signals than ventricular signals in both age groups, with overlapping 95 % confidence intervals between young and old mice.

### RT-qPCR

Age-associated changes in the expression of lysosomal/autophagy-related genes were analysed in young and old mouse hearts by normalising target gene expression against that of the housekeeping gene, *Gapdh*, (ΔCq) in R (v4.4.1). As shown in Figure 3, no significant differences (false discovery rate-adjusted *p* (*q*) > 0.5) in the expression of the lysosomal and autophagy genes *Atg12, Atp6v1a, Lamp2* and *M6pr*, or the antioxidant gene *Nfe2l2*, were identified. However, we noted increases in the expression of *Cd63* (*q* = 0.0756) and *Sqstm1* (*q* = 0.0018) in old hearts relative to young hearts. Full statistical outputs are provided in Supplementary Table S3.

**Figure 3.**
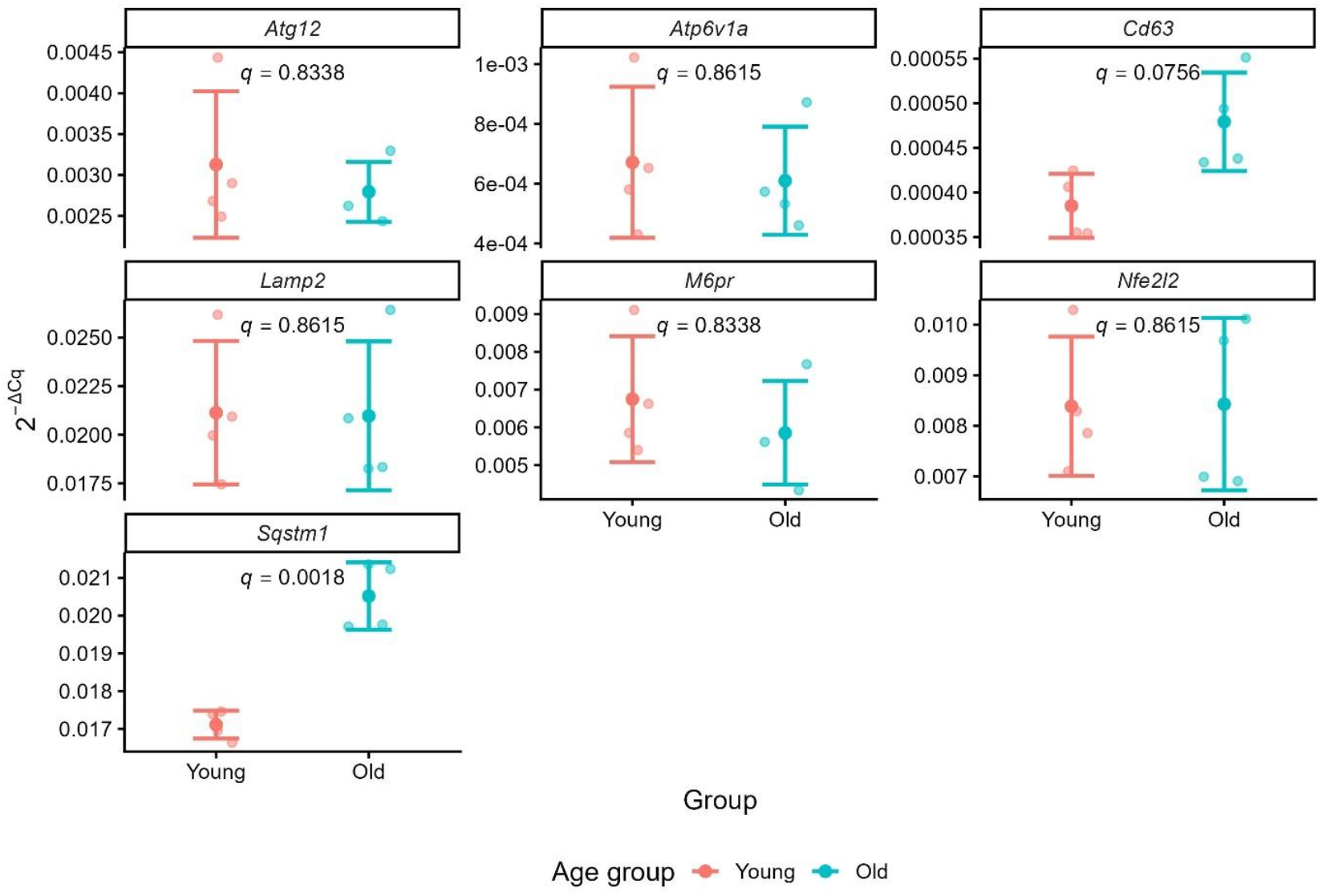
Gene expression in young and old mouse hearts (n = 4/group). Data represent transformed ΔCq values (2^−ΔCq^) calculated normalised to the housekeeping gene, *Gapdh*. Statistical comparisons performed using multiple unpaired *t-*tests with a two-stage Benjamini, Krieger and Yekutieli false discovery rate (FDR) correction. FDR-adjusted *p*-values (*q*-values) displayed on the graphs.

## Discussion

This study investigated age-associated changes in cardiac lysosomal properties using IVIS Lysotracker imaging together with RT-qPCR of selected lysosomal and autophagy-related genes. Our central hypothesis was that ageing would alter both the abundance of acidic organelles and the acidification machinery that supports them, leading to measurable differences in Lysotracker uptake and transcriptional profiles. Because Lysotracker accumulates in a protonated form within acidic compartments and cannot, on its own, resolve lysosomes from late endosomes, whole-organ signal is best interpreted as a proxy for the overall pool (number/volume) of acidic vesicles rather than absolute pH values (*27*). Using an optimised dual-sided 2D imaging strategy (∼5 s per orientation; ∼10 s total per heart), we observed no significant age-related differences in global Lysotracker fluorescence, indicating that the overall pool of acidic organelles is preserved in this aged group (18 months old hearts). This interpretation is reinforced by the stable expression of *Atp6v1a*, a catalytic v-ATPase subunit required for proton translocation and lysosomal acidification (*28*), and *Lamp2*, a lysosomal membrane protein that contributes to structural stability and chaperone-mediated autophagy (*5, 29*). Examining lysosome number in ageing hearts may be important to determine whether changes in quantity contribute to age-related decline, and the finding of stable numbers directs attention toward potential alterations in lysosomal function or efficiency instead.

The choice of fast 2D imaging was deliberate. While the IVIS Spectrum platform supports 3D fluorescence tomography (*30*), extended acquisition time risks *ex vivo* pH drift and probe signal decay. By imaging both anterior and posterior orientations, we maximised fluorescence capture while minimising artefacts, enabling confident assessment of whole-heart acidic-vesicle burden.

We also observed significantly higher Lysotracker signal in atrial regions compared to ventricles. This aligns with evidence that atrial cardiomyocytes are enriched in acidic vesicular stores, including atrial granules that interface with sarcoplasmic reticulum and mitochondria, supporting the atrium’s specialised role in Ca^2+^ handling and metabolic regulation (*31*). Complementing this, our group previously (*10*) demonstrated that lysosomal Ca^2+^ release pathways strongly influence atrial electrophysiology, linking acidic organelle signalling to rhythm control. Together, these studies validate the sensitivity of IVIS Lysotracker imaging in capturing physiologically meaningful spatial differences in acidic organelle distribution.

Although no overall age effect was detected, we noted batch-to-batch variability in Lysotracker signals. In Batch 2, fluorescence was slightly lower. Biological differences may also have contributed, since young mice in Batch 2 were 4 months old compared with 2 months in Batches 1 and 3. Importantly, imaging conditions were kept constant across experiments, and batch was included as a fixed effect in the statistical model to account for this variability. These measures ensure that our conclusion, that the acidic-vesicle pool and acidification capacity are maintained with ageing, is robust and not confounded by procedural or biological variability. A limitation of IVIS fluorescence imaging is its depth-dependent signal attenuation due to tissue absorption and scattering. Although the murine left ventricular wall thickness (∼1–1.5 mm) lies within the reported effective detection depth of IVIS (∼2–5 mm)(*32-34*), photon propagation is strongly surface-weighted, resulting in preferential detection of fluorophores located in superficial myocardial layers. Consequently, Lysotracker signal likely reflects predominantly epicardial cardiomyocytes rather than uniformly sampling the full transmural myocardium. IVIS measurements should therefore be interpreted as semi-quantitative and depth-biased, providing relative comparisons rather than absolute estimates of total myocardial lysosomal content. Furthermore, changes in tissue composition or wall thickness associated with disease may influence signal intensity independently of lysosomal biology. These considerations underscore that IVIS Lysotracker imaging is best suited for longitudinal or groupwise comparisons under similar imaging geometries, and complementary *ex vivo* or histological validation remains important for assessing transmural lysosomal distribution.

Our qPCR analysis revealed minimal transcriptional remodelling, in line with preserved whole-heart Lysotracker signal. However, as the assessment of autophagy-associated gene expression cannot measure post-translational modification and degradation of autophagy machinery, this approach may not fully capture the extent of cardiac autophagic function (*35*). Therefore, minor upregulation of *Sqstm1* (p62), a selective-autophagy adaptor central to proteostasis and oxidative stress responses, and *Cd63*, a marker of late endosomes and MVBs, may be indicative of enhanced autophagy activity (*36, 37*). *Cd63* upregulation may also reflect changes in vesicular trafficking and extracellular vesicle (EV) release, processes previously linked to ageing and intercellular signalling (*38, 39*). Given the stable expression of *Lamp2* and *Atp6v1a*, these changes are likely to be consistent with increased autophagy demand rather than impaired degradation (*5, 29, 40*), although functional assays and assessment of autophagic protein expression are required to support this. Furthermore, as Lysotracker preferentially labels highly acidic lysosomes, and late endosomes are less acidic, the lack of fluorescence increase potentially suggests that late endosomes (measured via *Cd63*) contribute minimally to the whole-heart signal (*27*).

Future studies should focus on three priorities. First, clarify organelle-specific contributions by combining IVIS imaging with ratiometric pH probes, compartment-specific labelling (e.g., LAMP2 vs. CD63), and V-ATPase inhibition assays to distinguish lysosomes from late endosomes (*27*). Second, resolve cell-type and regional drivers of transcriptional changes using spatial transcriptomics and single-nucleus RNA-seq, particularly in atrial regions where Lysotracker signals were highest. This builds on findings that atrial cardiomyocytes contain specialised acidic vesicular stores that influence electrophysiology via lysosomal Ca^2+^ signalling (*10, 31*). Finally, functional assays measuring autophagic flux, cathepsin activity, and extracellular vesicle release should determine whether subtle age-associated changes in *Sqstm1* expression reflects increased autophagic function or impaired clearance (*29, 40*). Together, these approaches will define the mechanisms linking lysosomal Ca^2+^ signalling and atrial specialisation to cardiac ageing and age-related cardiac diseases such as AF, informing precision strategies to maintain lysosomal homeostasis.

## Conclusion

Taken together, these findings support a model in which the abundance of acidic vesicles and their acidification capacity are maintained with age (IVIS imaging results - *Atp6v1a, Lamp2*, figure 3), it is possible that lysosome-associated regulatory pathways may undergo adaptive remodelling but this needs further validation and exploration. Physiologically, these adaptations occur alongside atrial enrichment of acidic vesicular stores (*31*) and lysosomal Ca^2+^ signalling contributions to atrial excitability (*10*), highlighting the dynamic regulation of lysosome-associated pathways during cardiac ageing. However, we cannot say with certainty the true depth imaged by IVIS lysotracker imaging in relation to its optical penetration., while bulk RT-qPCR averages expression across multiple cardiac cell types may not adequately reflect autophagic function. Consequently, subtle or cell-type–specific alterations in lysosomal function may have gone undetected. The close spatial proximity of lysosomes to the sarcoplasmic reticulum in myocytes(*10, 16*), highlights a potential structural and functional coupling that may influence intracellular Ca^2+^ dynamics. Investigating the quantity of acidic vesicles and their acidification capacity will provide critical insight into how lysosomal dysfunctions contribute to disease progression as well as in aging. Future studies employing older mice (e.g. 24 months or older), higher-resolution imaging, LAMP2 immunostaining, and assays of lysosomal enzyme activity will be important to resolve regional and cellular differences.

## Limitations

This study has several limitations. First, IVIS fluorescence imaging is inherently depth-limited and surface-weighted. Because both excitation and emission light are strongly attenuated in cardiac tissue, the Lysotracker signal predominantly reflects the epicardial layer, and potential age-related alterations within deeper myocardial regions may have gone undetected.

Second, Lysotracker fluorescence reflects lysosomal acidity rather than degradative function or total lysosome number. Thus, changes in lysosomal pH or enzyme activity could occur without corresponding differences in fluorescence intensity, limiting interpretation of lysosomal function.

Third, bulk RT-qPCR analysis of *Lamp2* provides an averaged measure of gene expression across heterogeneous cardiac cell types. Consequently, distinct transcriptional alterations within specific cardiac cell populations, such as cardiomyocytes, fibroblasts, or endothelial cells, is not resolved.

Fourth, the use of *Lamp2* as a single lysosomal marker does not capture the full scope of lysosomal biogenesis or degradative capacity. Inclusion of additional lysosomal or autophagy-related genes (e.g., *Lamp1, Ctsd, Tfeb*) and protein or activity-based assays would yield a more comprehensive assessment.

Finally, all imaging and molecular analyses were conducted *ex vivo*, where altered perfusion, pH, and metabolic state may influence Lysotracker uptake and retention. Future studies integrating confocal or super-resolution imaging, immunostaining for multiple lysosomal markers, enzyme activity assays, and single-cell or spatial transcriptomics would help delineate cell-type, specific and functional lysosomal remodelling during cardiac ageing.

## Acknowledgements

This study was supported by the Ellis T Davies Fellowship Endowment, University of Liverpool (PI RABB). GRM acknowledges support from the BBSRC (grant number BB/V01840X/1). For the purpose of Open Access, the author has applied a CC BY public copyright license to any Author Accepted Manuscript version arising from this submission. We thank Katharine Gittins and Bradley Westley from the BMS for support in animal work. We thank Prof Ian Copple for access to laboratory equipment for running the RT-qPCR experiments.

## Authorship contribution statement

RABB: Conceptualization and reviewing; RABB: Fund acquisition. RABB, JA, HC, MM: Investigation, writing the first draft, reviewing and editing. MM and MM: IVIS technical development. NH, PS, HP: formal analysis, writing, reviewing and editing. HC, FP, JH, EA, LH, OA: Serum, tissue collection and RT qPCR studies. PS and GM: analysis and scripts.

## Data availability

Files have been uploaded onto figshare [https://doi.org/10.6084/m9.figshare.29984677]. The corresponding author can be contacted for any further details.

## Methods

### IVIS Imaging

#### Ethical approval & animal details

All animal procedures were conducted in accordance with institutional and national ethical guidelines and were approved by [University of Liverpool, Animal Welfare and Ethical Review Body (AWERB)]. Mice were euthanized using rising concentrations of carbon dioxide (CO_2_), in accordance with Schedule 1 methods as defined by the UK Animals (Scientific Procedures) Act 1986 and the Home Office guidelines. Hearts from 14 male C57BL/6J mice (2–4-month-old or 18-month-old; n = 7 per age group) were included in the fluorescence imaging analysis. A total of 16 animals were used across three imaging batches (purchased from Janvier-Labs and Charles River Laboratories), but two samples were excluded: one old mouse (O4) was treated with buffer only (negative control), and one young mouse (Y4), assigned as a second control. The buffer-only heart served to confirm successful dye loading and IVIS instrument performance; thus, exclusion of the second control was not expected to affect data integrity or interpretation. An overview of batch composition is shown in Table 1. Corresponding individual animal body and heart weights are provided in Supplementary Table S4.

**Table 1.** Batch Structure and Control. Details of batch dates, group sizes, and included samples for the IVIS experiment. Y = young; O = old.

| Batch | Date | Group Sizes | Notes |
| --- | --- | --- | --- |
| 1 | 13 MAY 2025 | 3 old/ 3 young | Two excluded. Final (Y1-Y2, O1-O2, Control) |
| 2 | 7 JUL 2025 | 3 old / 3 young | All included (Y3–Y5, O3–O5) |
| 3 | 4 AUG 2025 | 2 old / 2 young | All included (Y6–Y7, O6–O7) |

#### Reagent Preparation

**LysoTracker™ Red DND-99** (Thermo Fisher Scientific, Cat. No. L7528; 1 mM stock) was diluted by adding 0.5 μL to 5 mL of sterile, ice-cold Balanced Phosphate Solution (BPS) and protected from light.

**Tyrode’s solution** (composition in Table 2) was thawed from pre-prepared aliquots stored at –20 °C and supplemented with 0.1 mL heparin per ∼40 mL total volume immediately before use. This solution was used for retrograde perfusion during heart preparation.

**Table 2.** Composition of Tyrode’s Solution. Composition of Tyrode’s solution showing concentrations (mM) of each component, adjusted to pH 7.4 with NaOH.

| Component | Amount (mM) |
| --- | --- |
| KCl | 5 |
| NaCl | 135 |
| NaH<sub>2</sub>PO<sub>4</sub> | 0.33 |
| Na-pyruvate | 5 |
| HEPES | 10 |
| Mannitol | 15 |
| Glucose | 5 |
| CaCl<sub>2</sub> | 2 |
| MgCl<sub>2</sub> | 1 |

**Phosphate Buffer Saline** (ice-cold, Gibco #10010-015) was used for control treatment in Batch 1.

#### Phantom Studies

A preliminary phantom experiment was conducted to validate excitation/emission filter settings and optimize fluorescence detection before mouse imaging. LysoTracker™ Red DND-99 (Thermo Fisher Scientific, Cat. No. L7528; 1 mM stock) and LysoTracker™ Green (Thermo Fisher Scientific, Cat. No. 497269; 1 mM stock) were each diluted in 5 mL of sterile, ice-cold Balanced Phosphate Solution (PBS) and kept on ice, protected from light. Samples were transferred to a black-bottom 96-well plate to minimize background fluorescence, reduce light scattering, and prevent signal saturation. Several excitation/emission filter combinations were tested during the dry run, and based on these results and manufacturer specifications, the 570/620 nm filter set was selected for all subsequent experiments. This setting closely approximates the LysoTracker™ Red excitation/emission maxima (577/590 nm) and provided optimal signal quality confirmed using phantom dye controls.

#### Heart Collection, Imaging, and Post-Processing

Mice were euthanized in pairs to standardize timing. Hearts were rapidly dissected, weighed, and placed in ice-cold Tyrode’s solution containing heparin. Blood was collected immediately into Microvette® 500 CAT tubes (SARSTEDT, Cat. No. 20.1343), allowed to clot for 30 min at room temperature, and centrifuged at 3,000 rpm for 15 min at 20 °C. Serum supernatants were snap-frozen in liquid nitrogen and stored at −80 °C for subsequent analyses.

Epicardial fat on the heart surface was gently trimmed to expose the aorta, which was retrogradely perfused with Tyrode’s solution to remove residual blood. LysoTracker™ Red was then delivered via the aorta using a 1 mL syringe and plastic cannula, followed by 20 min incubation on ice, protected from light. Hearts were imaged on the IVIS® Spectrum system (PerkinElmer) with the chamber maintained at 28–30 °C, using the 570/620 nm filter set. Each heart was imaged in both anterior (front) and posterior (flipped) orientations (Figure 4).

**Figure 4.**
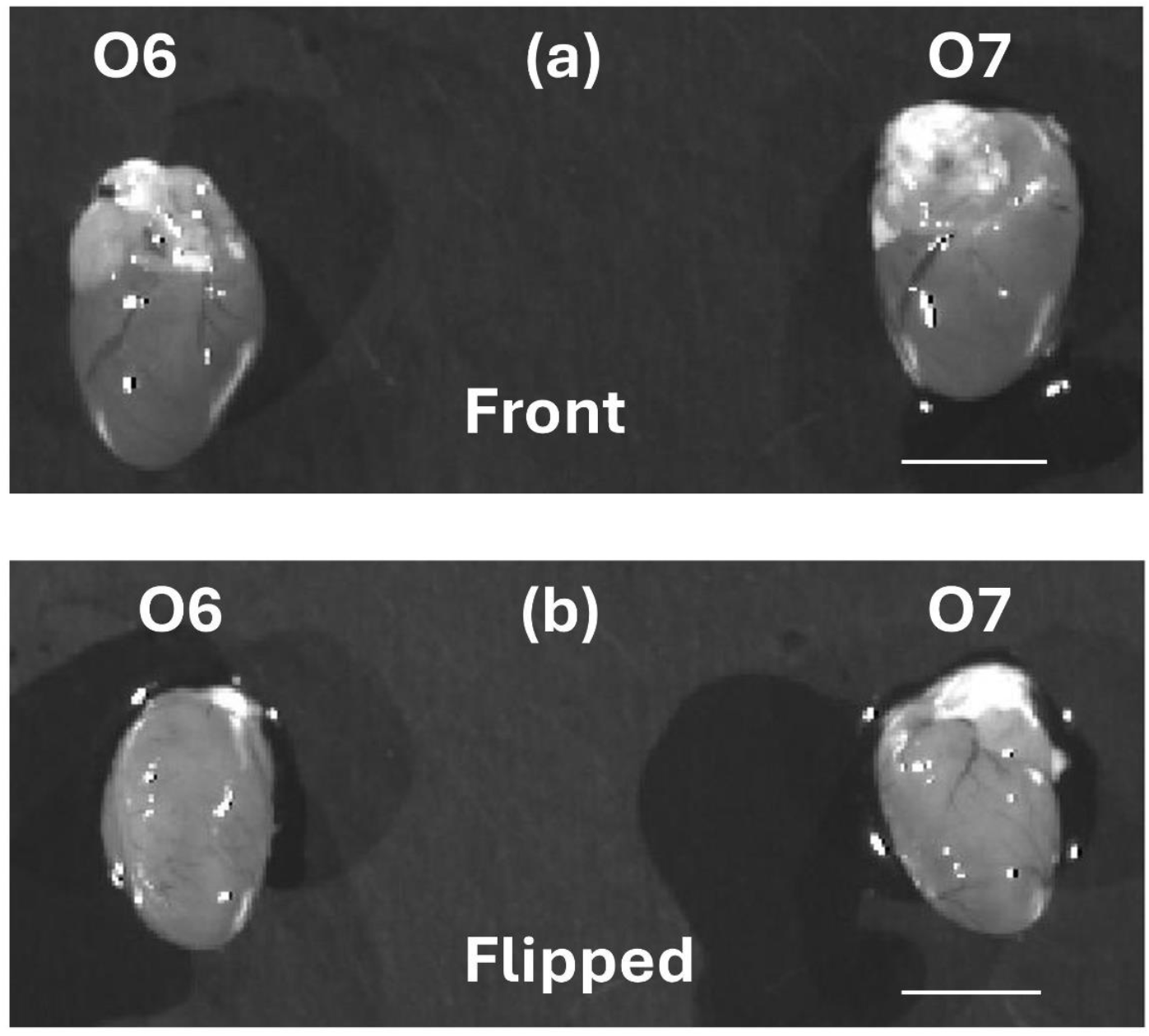
Orientation of mouse hearts during IVIS imaging. Representative image showing mouse hearts positioned in the front (anterior) and flipped (posterior) orientation within the IVIS® Spectrum imaging chamber for fluorescence acquisition. O = old animal. Scale bar 0.5 cm.

After imaging, hearts were snap-frozen in liquid nitrogen and stored at −80 °C for downstream qPCR analysis of selected lysosomal, late endosomal, and autophagy-related markers to complement IVIS measurements and evaluate age-related changes in lysosomal function and autophagy flux. Considering that LysoTracker™ Red accumulates in acidic intracellular compartments without distinguishing between lysosomes and late endosomes, signals were interpreted as a general proxy for acidic organelle abundance rather than a direct measure of lysosomal activity (*27*).

#### Image Analysis

Regions of interest (ROIs) were manually drawn using the Living Image® v4.5.4 software, guided by anatomical landmarks with expert input. For each heart, ROIs for whole heart, atrium, and ventricle were defined in both anterior and posterior orientations; paired orientation values were summed to yield one value per anatomical region per mouse. Fluorescence was quantified as Average Radiant Efficiency [p/s/cm^2^/sr]/[µW/cm^2^], enabling direct comparison across ROIs of different sizes. A representative example of ROI placement and signal distribution is shown in Figure 5.

**Figure 5.**
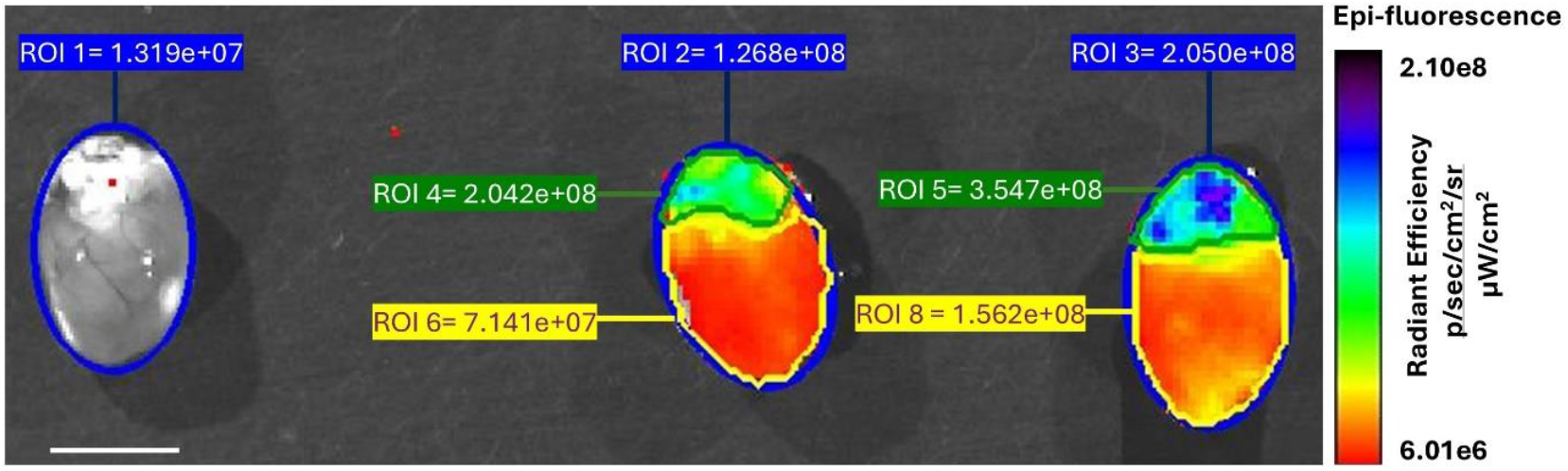
Representative IVIS® Spectrum image from Batch 1 (Control, O1, O2; anterior view). Manually defined ROIs and corresponding fluorescence values. Blue = whole heart, yellow = ventricle, green = atrium. Values represent Average Radiant Efficiency [p/s/cm^2^/sr]/[µW/cm^2^]. Scale bar 0.5 cm.

#### Data Analysis

Fluorescence signals (Average Radiant Efficiency) were exported from Living Image® (v4.7.3) and analyzed in R (v4.4.1). Anterior and posterior ROI signal intensity values for each region were summed to capture total signal (mitigating superficial bias inherent to IVIS imaging) and to minimize orientation-related variability. Linear mixed-effects models (lme4) included Age (young vs. old), Region (atrium vs. ventricle), and Batch were used as fixed effects, with Mouse ID as a random intercept. Both raw and log_10_-transformed data were tested; diagnostics supported the log_10_ model, which better satisfied assumptions. Estimated marginal means (EMMs) with 95 % CIs were computed using the *emmeans* R package and back-transformed for interpretation. Final conclusions were based on the log_10_-transformed, batch-adjusted model.

### RT-qPCR Analysis

RNA was isolated from mouse hearts using the Monarch® Spin RNA Isolation Kit (NEB, T2010) according to the manufacturer’s protocol and RNA concentration and purity was assessed using a NanoDrop™ ND-1000 spectrophotometer. Complementary DNA (cDNA) was synthesised from RNA using the LunaScript® RT SuperMix Kit (NEB, E3010) and the following thermal cycling conditions: 2 mins at 25 °C, 10 mins at 55 °C, then 1 min at 95 °C. qPCR reactions were prepared using the Luna® Universal qPCR Master Mix (NEB, M3003) and run on a QuantStudio ™ 7 Pro real-time PCR system in technical duplicates under the following thermal cycling conditions: 60 s at 95 °C, then 40 cycles of 15 s at 95 °C followed by 30 s at 60 °C. Following amplification, a melt curve was generated across a temperature range of 65–95 °C.

### Primer design and validation

Primers were designed for *Lamp2, Cd63, M6pR, Atpv1a, Gapdh, Atg12, Nfe2l2*, and *Sqstm1* using NCBI Primer-BLAST, targeting exon–exon junctions to avoid genomic DNA amplification. Amplicon sizes (70–200 bp), GC content (40–60 %), and melting temperatures (∼60 °C) were optimized, while avoiding self-dimers, secondary structures, and primer–dimers (Supplementary Table S5). For this purpose, RNA was extracted from a young mouse sample (Y0, excluded from the imaging study) and a cDNA template standard curve ranging from 0-50 ng/μL was generated to validate primer efficiency. Acceptance criteria were defined as R^2^ ≥ 0.98, slope –3.1 to –3.6, and efficiency 90–110 %. All primers met these thresholds except for *Atp6v1a* and (R^2^ = 0.969) and *Cd63* (<u>R^2^ = 0.978</u>); however, these primers were accepted as these values were close to the defined threshold and the slope and efficiencies for these primers met the acceptance criteria (Supplementary Table S5).

### qPCR setup and controls

RNA and cDNA samples and qPCR reactions were prepared as previously described. RNA concentrations were standardised to 15 ng/μL. Non-template controls (NTC; water + master mix) and no-RT controls (NRT; RNA + water + no RT mix) were included in each run to exclude reagent contamination and genomic DNA carryover. All data were processed in R (v4.4.1). Target gene expression was normalised to that of the housekeeping gene, *Gapdh* and the effect of ageing on gene expression was assessed statistically in GraphPad Prism 10.2.3 using multiple unpaired *t-*tests with a two-stage Benjamini, Krieger and Yekutieli false discovery rate (FDR) correction.

All raw IVIS imaging data, qPCR measurements, and R analysis scripts supporting this study are openly available on Figshare at “https://doi.org/10.6084/m9.figshare.29984677 “.

## SUPPLEMENTARY MATERIAL

**Supplementary Figure S1.**
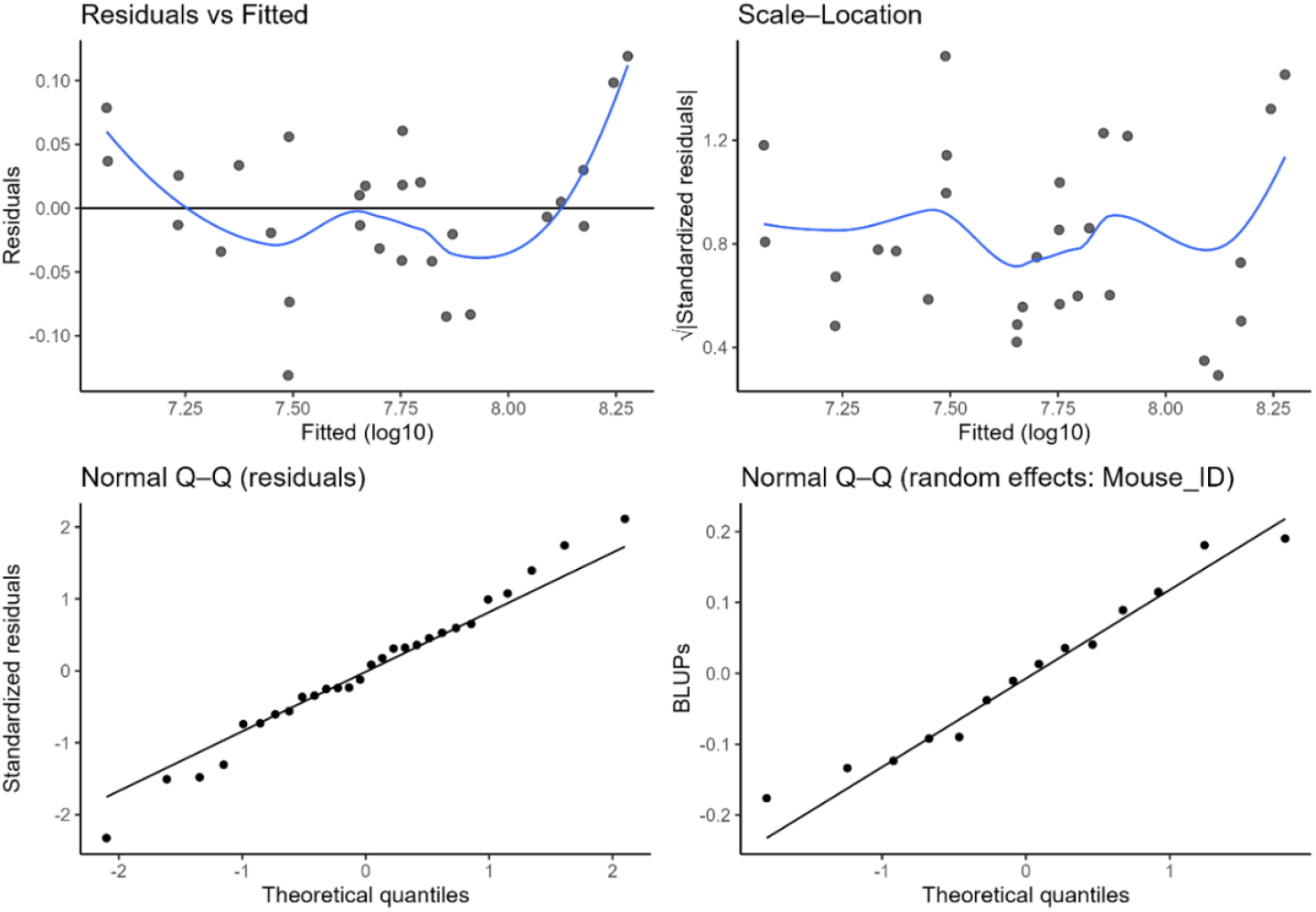
Plots for the linear mixed-effects model assessing the effects of Age group, Region, and Batch on log10_{10}10-transformed IVIS signals. **Top left:** Residuals vs. fitted values, showing no strong non-linear patterns but slight curvature, suggesting minor model misfit. **Top right:** Scale–location plot, indicating roughly constant variance across fitted values, with some spread differences at the extremes. **Bottom left:** Normal Q–Q plot of residuals, showing approximate normality with minor deviations in the tails. **Bottom right:** Normal Q– Q plot for random effects (Mouse_ID), indicating that random intercepts are approximately normally distributed. These diagnostics support the appropriateness of the model for the data, with no major violations of assumptions.

**Supplementary Table S1.** Estimated marginal means (EMMs) ± 95% confidence intervals for each Age × Region combination from the linear mixed-effects model, with both log10_{10}10-transformed and back-transformed (geometric mean) signal values. Back-transformed values are presented in photons/sec/cm^2^/sr for direct interpretability.

| Region | Age_Group | Mean_log10 | Lower95_log10 | Upper95_log10 | GeoMean | Lower95 | Upper95 |
| --- | --- | --- | --- | --- | --- | --- | --- |
| Atrium | Young | 7.927 | 7.796 | 8.059 | 84500000 | 62500000 | 115000000 |
| Atrium | Old | 7.977 | 7.846 | 8.108 | 94800000 | 70100000 | 128000000 |
| Ventricle | Young | 7.506 | 7.375 | 7.637 | 32100000 | 23700000 | 43400000 |
| Ventricle | Old | 7.556 | 7.424 | 7.687 | 36000000 | 26500000 | 48600000 |

**Supplementary Table S2.** Full ANOVA results from the linear mixed-effects model testing the effects of Age group, Region, and Batch on log10_{10}10-transformed IVIS signals, with Mouse ID as a random intercept. Values include degrees of freedom (df), F-statistics, and *p*-values for each fixed effect.

| Effect | Sum Sq | Mean Sq | NumDF | DenDF | F value | Pr(>F) |
| --- | --- | --- | --- | --- | --- | --- |
| Age_Group | 0.0023 | 0.0023 | 1 | 10 | 0.3767 | 0.5531 |
| Region | 1.2429 | 1.2429 | 1 | 13 | 207.3086 | 0 |
| Batch | 0.2075 | 0.1037 | 2 | 10 | 17.3036 | 0.0006 |

**Supplementary Table S3.** Summary of multiple unpaired *t*-tests and Benjamini, Krieger and Yekutieli FDR correction testing the effects of age group ΔCq values measured by RT-qPCR. Values include unadjusted *p*-values, test statistics (t ratio), degrees of freedom (DF), FDR-adjusted *p*-values (*q*-value) for each gene.

|  | **P value** | **t ratio** | **DF** | **q value** |
| --- | --- | --- | --- | --- |
| **Atg12** | 0.5504 | 0.6325 | 6 | 0.8338 |
| **Atp6v1a** | 0.7484 | 0.3359 | 6 | 0.8615 |
| **Cd63** | 0.0250 | 2.970 | 6 | 0.0756 |
| **Lamp2** | 0.9470 | 0.06935 | 6 | 0.8615 |
| **M6pr** | 0.4225 | 0.8606 | 6 | 0.8338 |
| **Nfe2l2** | 0.9951 | 0.006422 | 6 | 0.8615 |
| **Sqstm1** | 0.0003 | 7.451 | 6 | 0.0018 |

**Supplementary Table S4.** Body and heart weights of all mice used in IVIS experiments. Animal weight and heart weight were recorded immediately after sacrifice for each mouse in the Old (O1–O7) and Young (Y1–Y7) groups, with one Control sample included. Values are presented in grams (g).

| **Mouse ID** | **Animal Weight (g)** | **Heart Weight (g)** |
| --- | --- | --- |
| O1 | 29.4 | 0.20 |
| O2 | 33.6 | 0.27 |
| O3 | 36.78 | 0.36 |
| O4 | 36.04 | 0.26 |
| O5 | 36.59 | 0.26 |
| O6 | 31.7 | 0.25 |
| O7 | 33.6 | 0.30 |
| Y1 | 25.8 | 0.22 |
| Y2 | 21.8 | 0.28 |
| Y3 | 31.64 | 0.26 |
| Y4 | 30.56 | 0.39 |
| Y5 | 30.48 | 0.26 |
| Y6 | 20.19 | 0.18 |
| Y7 | 25.69 | 0.18 |
| Control | 32.8 | 0.33 |

**Supplementary Table S5.**
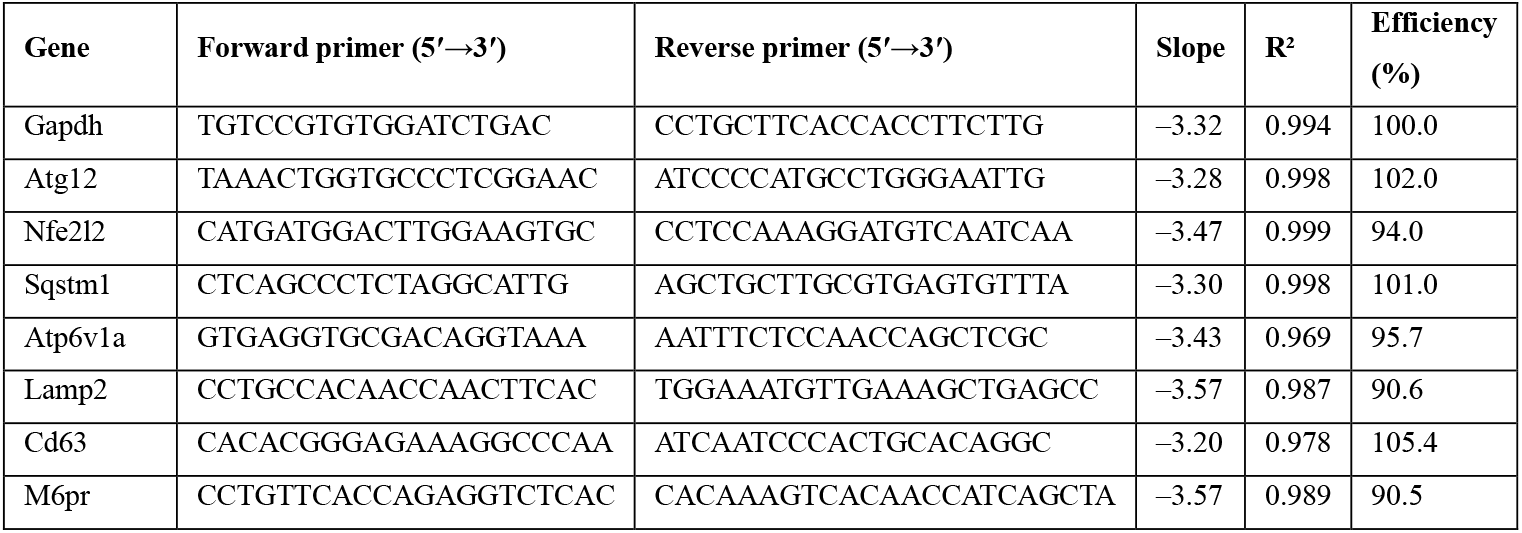
Validated primer sequences and performance parameters used for qPCR. All primers demonstrated acceptable qPCR performance, with slopes between –3.1 and –3.6 and efficiencies between 90–110 %. R^2^ values were ≥0.98 for all assays, with the exception of *Atp6v1a* (R^2^ = 0.969) and *Cd63* (R^2^ = 0.978), which fell just below the R^2^ cut off but retained excellent efficiency, supporting their suitability for use.

